# The effect of root exudates on the transcriptome of rhizosphere *Pseudomonas* spp

**DOI:** 10.1101/2021.01.08.425997

**Authors:** Olga V. Mavrodi, Janiece R. McWilliams, Jacob O. Peter, Anna Berim, Karl A. Hassan, Liam D. H. Elbourne, Melissa K. LeTourneau, David R. Gang, Ian T. Paulsen, David M. Weller, Linda S. Thomashow, Alex S. Flynt, Dmitri V. Mavrodi

## Abstract

Plants live in association with microorganisms that positively influence plant development, vigor, and fitness in response to pathogens and abiotic stressors. The bulk of the plant microbiome is concentrated belowground at the plant root-soil interface. Plant roots secrete carbon-rich rhizodeposits containing primary and secondary low molecular-weight metabolites, lysates, and mucilages. These exudates provide nutrients for soil microorganisms and modulate their affinity to host plants, but molecular details of this process are largely unresolved. We addressed this gap by focusing on the molecular dialogue between eight well-characterized beneficial strains of the *Pseudomonas fluorescens* group and *Brachypodium distachyon*, a model for economically important food, feed, forage, and biomass crops of the grass family. We collected and analyzed root exudates of *B. distachyon* and demonstrated the presence of multiple carbohydrates, amino acids, organic acids and phenolic compounds. The subsequent screening of bacteria by Biolog Phenotype MicroArrays revealed that many of these metabolites provide carbon and energy for the *Pseudomonas* strains. RNA-seq profiling of bacterial cultures amended with root exudates revealed changes in the expression of genes encoding numerous catabolic and anabolic enzymes, transporters, transcriptional regulators, stress response, and conserved hypothetical proteins. Almost half of the differentially expressed genes mapped to the variable part of the strains’ pangenome, reflecting the importance of the variable gene content in the adaptation of *P. fluorescens* to the rhizosphere lifestyle. Our results collectively reveal the diversity of cellular pathways and physiological responses underlying the establishment of mutualistic interactions between these beneficial rhizobacteria and their plant hosts.

## INTRODUCTION

Plants are meta-organisms or holobionts that rely in part on their microbiome for specific functions and traits. The ability of the plant microbiome to influence plant development, vigor, health and fitness in response to abiotic stressors associated with global climate change is documented by numerous studies (Lugtenberg and Kamilova, 2009). There is mounting evidence that plants actively recruit beneficial microbiomes, but many aspects of this process are still very much a black box (Reinhold-Hurek et al., 2015). The foundation for the differential affinity of rhizobacteria towards host plants is built upon complex chemical cross talk between microorganisms and plant roots. Up to 40% of photosynthetically fixed carbon is released by plant roots in the form of exudates and secretions, lysates, and mucilages (Curl and Truelove, 1986; Lynch, 1990; Whipps, 1990; Badri and Vivanco, 2009). The release of these compounds is actively controlled in response to environmental stimuli, and the composition of rhizodeposits varies greatly according to plant species and physiological condition (Lynch, 1990; Nguyen, 2003; Phillips et al., 2004; De-la-Pena et al., 2008). The presence and composition of exudates strongly impact soil microorganisms, which is consistent with the idea that plants actively select and shape their root microbiota (Zolla et al., 2013).

Primary root exudates include simple and complex sugars, amino acids, polypeptides and proteins, organic, aliphatic and fatty acids, sterols and phenolics (Nguyen, 2003; Badri and Vivanco, 2009; Badri et al., 2009). These compounds serve as carbon and energy sources for rhizobacteria, and the presence of the intact corresponding catabolic pathways is essential for competitive colonization of roots and disease suppression (Lugtenberg et al., 2001; Kamilova et al., 2005; Lugtenberg and Kamilova, 2009). Root exudates also contain numerous signal molecules and secondary metabolites, the significance of which is only now emerging (Walker et al., 2003; Bais et al., 2005; Bais et al., 2006). A handful of analyses of plant-induced gene expression by transcriptional profiling *in vitro* (Mark et al., 2005) or in the rhizosphere (Silby and Levy, 2004; Ramos-Gonzalez et al., 2005; Matilla et al., 2007; Barret et al., 2009) have identified multiple genes that are differentially regulated by exposure to roots or root exudates. Bacterial pathways expressed during rhizosphere colonization control utilization of plant-derived metabolites (Simons et al., 1996; Simons et al., 1997; Camacho-Carvajal, 2001; Lugtenberg and Kamilova, 2009), motility and chemotaxis (de Weert et al., 2002; Lugtenberg and Kamilova, 2009), phase variation (Dekkers et al., 1998; Sanchez-Contreras et al., 2002; van den Broek et al., 2005), outer membrane integrity (de Weert et al., 2006; Lugtenberg and Kamilova, 2009), and the ability to sequester limiting resources (Raaijmakers et al., 1995) and resist environmental stresses (Sarniguet et al., 1995; Miller and Wood, 1996; van Veen et al., 1997; Schnider-Keel et al., 2001). In its spatial and temporal properties, root colonization resembles biofilm formation, and biofilm-related pathways also have been implicated in adhesion to seeds and roots and rhizosphere colonization (Espinosa-Urgel et al., 2000; Hinsa et al., 2003; Yousef-Coronado et al., 2008; Fuqua, 2010; Martinez-Gil et al., 2010; Nielsen et al., 2011; Zboralski and Filion, 2020). Finally, root exudates strongly affect the expression of diverse plant growth promotion and biocontrol genes (Vacheron et al., 2013). Over the past decade, the genomes of numerous rhizosphere strains have been sequenced and analyzed, but functional genomics studies of rhizosphere competence lag behind the availability of sequence data.

This study explored the molecular dialogue between the model host plant *Brachypodium distachyon* and several well-characterized rhizosphere strains of the *Pseudomonas fluorescens* group. *Brachypodium* is a small annual grass originating in semi-arid regions of the Middle East that has emerged as a prime model for economically important food, feed, forage and biomass crops of the grass family (Bevan et al., 2010; Brkljacic et al., 2011). *Brachypodium* has small stature, a compact genome, short generation time, and has served for phenotype investigations into natural variations in morphology, growth habit, stem length, stem density, and cell wall composition, seed proteome characteristics, codification and description of growth stages, and natural variation in flowering time in response to vernalization (Schwartz et al., 2010; Hong et al., 2011; Tyler et al., 2014). The biology, extensive collection of resources, and research tools make *B. distachyon* an attractive model to investigate interactions between plants and root-associated microbes. Pseudomonads are ubiquitous Gram-negative γ-proteobacteria known for their utilization of numerous organic compounds as energy sources, production of diverse secondary metabolites and resistance to antimicrobials. These bacteria can colonize eukaryotic hosts and include both commensals and economically important pathogens of plants and animals (Moore et al., 2006; Schroth et al., 2006; Yahr and Parsek, 2006). The genus *Pseudomonas* currently comprises >100 named species that have been separated based on multilocus sequence analysis into 14 species groups (Garrido-Sanz et al., 2016; Hesse et al., 2018). The *P. fluorescens* group is the most diverse regarding both the genetic distances within it, the number of species and the large pan-genome that makes up >50% of the pan-genome of the genus as a whole (Loper et al., 2012). The group also encompasses an unusually high proportion of strains that inhabit the plant rhizosphere and possess plant growth promoting and biocontrol properties. We focused on eight well-studied strains of the *P. fluorescens* complex that are supported by years of studies, numerous refereed publications and high-quality genome sequences. By profiling transcriptomes of these strains during growth in root exudates of *B. distachyon*, we revealed the diversity of cellular pathways and physiological responses that underlie the establishment of mutualistic interactions between beneficial rhizobacteria and the host plant. Our results also confirmed that root exudates contain carbohydrates, amino acids, organic acids and phenolics that serve as carbon and energy sources for rhizobacteria. The rhizodeposits also contained osmoprotectants that may help microorganisms to persist in the rhizosphere of drought-stressed plants. The diversity of microbial genes perturbed by root exudates reflects the importance of the variable genome in adaptation of individual strains of *Pseudomonas* to the rhizosphere lifestyle.

## MATERIALS AND METHODS

### Bacterial Strains Used in the Study

The eight *Pseudomonas* strains used for this study are *P. synxantha* 2-79 (Thomashow and Weller, 1988), *P. fluorescens* SBW25 (Silby et al., 2009), *Pseudomonas* sp. R1-43-08 (Parejko et al., 2012), *P. brassicacearum* Q8r1-96 (Raaijmakers and Weller, 1998), *P. fluorescens* Q2-87 (Bangera and Thomashow, 1996), *P. chlororaphis* 30-84 (Thomashow et al., 1990), *P. fluorescens* Pf0-1 (Silby et al., 2009), and *P. protegens* Pf-5 (Howell and Stipanovic, 1980). The selected organisms have been studied extensively for their role in biological control and plant growth promotion (**Table S1**). The strains were maintained in the laboratory as frozen stocks (−80°C) and routinely cultured on King’s medium B (King et al., 1954).

### Propagation of Plants and Collection of Root Exudates

*B. distachyon* Bd21 was established from seed material obtained from the USDA-ARS Plant Germplasm Introduction and Testing Research Unit (Pullman, WA). *Brachypodium* seeds were imbibed for three days at 4°C and sown in 7×7 cm pots filled with Sunshine Potting Mix #4 (Sun Gro Horticulture, Agawam, MA). Plants were grown in an IR-89X (Percival Scientific, Perry, IA) controlled environment chamber retrofitted with 6500K and 3000K T5 54W grow lights (Spectralux) under a 20-h light, 24°C/4-h dark, 18°C cycle. Plants were watered and fertilized with Jack’s Professional Water-Soluble Fertilizer (20:20:20) (JR Peters, Allentown, PA). After 12 weeks and plant senescence, seeds were collected, processed, and stored under desiccant and dark conditions at room temperature.

To collect root exudates, seeds of *B. distachyon* Bd21 were surface-sterilized, pre-germinated, and placed in sterile 1 L wide-mouth glass jars containing 113 g of 6-mm glass beads and 25 mL distilled water. Jars were covered with vented caps and plants were grown hydroponically in an environmental controlled growth chamber under conditions described above. After 6 days, root exudates were extracted from individual jars and their sterility was confirmed by spotting on nutrient agar. Multiple batches of root exudates were collected, filtered (0.22 µm), aliquoted in Falcon tubes, lyophilized and stored at −80°C.

### Metabolomic Profiling of Root Exudates

Exudates were analyzed for primary metabolites at the Murdock Metabolomics Laboratory at Washington State University (Pullman, WA). Freeze-dried residues were suspended in 500 μL 50% aqueous acetonitrile and clarified by centrifugation for 20 min at 21,000× *g* and 4°C. The liquid chromatography mass spectrometry analysis was conducted with a Synapt G2-S quadrupole-ion mobility spectrometry-time of flight mass spectrometer system equipped with an Acquity ultra-performance liquid chromatograph (UPLC) and an Acquity photodiode array detector (all from Waters, Milford, MA). The exudate metabolites were separated on a SeQuant ZIC-pHILIC HPLC column (2.1 × 100 mm, 3 μm) (Milllipore Sigma, Burlington, MA) using acetonitrile with 0.1% formic acid as solvent B and water with 0.1% formic acid as solvent A at a flow rate of 400 μL min^−1^ and the following linear gradient extending over 14 min: 0 min, 80% B; 4 min, 80% B, 6min: 10% B; 7.5 min, 10% B; 10 min, 80% B; 14 min, 80% B. Mass spectra were collected in positive ion mode over a range of *m/z* 50-1200 with a scan time of 0.2 sec^−1^. The Q-TOF-MS source was at 3.0 kV and 120°C; the sampling cone at 40V, desolvation temperature was 250°C; cone gas and desolvation gas flow were at 0 and 850 L h^−1^, respectively. Leucine enkephalin was used for post-acquisition mass correction. Target compounds were visualized using selected ion chromatograms at 0.05 Da window width. The compound identification was based on comparison of chromatographic behavior and accurate masses to those of authentic standards.

For gas chromatography, derivatization was carried out using a modification of the procedure of Lee & Fiehn (Lee and Fiehn, 2008). The freeze-dried residues were suspended in 950 μL aqueous methanol (84% v/v) and clarified by centrifugation for 15 min at 21,000 × *g* at 4°C. The supernatants were spiked with 1 μg of the internal standard salicylic acid-d_6_ (C/D/N Isotopes, Quebec, Canada) and dried *in vacuo*. The dry residues were suspended in 10 μL of *O*-methoxylamine hydrochloride (30 mg mL^−1^ in anhydrous pyridine, both from Milllipore Sigma) and incubated while mixing (1000 RPM) for 90 min at 30°C. Subsequently, samples were derivatized with 90 μL of MSTFA with 1% TMCS (Thermo Fisher Scientific, Waltham, MA) for 30 min at 37°C. Gas chromatography-mass spectroscopy analysis was performed using a Pegasus 4D time-of-flight mass spectrometer (LECO, Saint Joseph MI) equipped with a MPS2 autosampler (Gerstel, Linthicum, MD) and a 7890A oven (Agilent Technologies, Santa Clara, CA). The derivatization products were separated on a 30-m, 0.25 mm i.d., 0.25 μm d_f_ Rxi-5Sil column (Restek, Bellefonte, PA) with an IntegraGuard pre-column using ultrapure He at a constant flow of 0.9 mL min^−1^ as carrier gas. The linear thermal gradient started with a one-minute hold at 70°C, followed by a ramp to 300°C at 10°C min^−1^. The final temperature was held for 5 min prior to returning to initial conditions. Mass spectra were collected at 17 spectra sec^−1^. Peak identification was conducted using the Fiehn primary metabolite library (Kind et al., 2009) and an identity score cutoff of 700. Additionally, authentic standards for a number of primary metabolites were analyzed under identical conditions and the data used to compare the chromatographic behavior. Peak alignment and spectrum comparisons were carried out using the Statistical Compare feature of ChromaTOF software (LECO).

### Isolation of RNA and RNA-Seq

The strains were pre-grown overnight at 25°C on 21C agar (Halverson and Firestone, 2000) with 10 mM glucose and then subcultured into 96-well microplates containing liquid 21C-glucose medium amended with 20-fold concentrated *Brachypodium* exudates. The control cultures were grown under identical conditions in the absence of exudates. All treatments were inoculated at OD_600_ of 0.1 and incubated until cultures entered mid-exponential growth phase at 25°C in an atmosphere of 15% oxygen (created by a ProOx P110 oxygen controller (BioSpherix, Parish, NY) with a hypoxia C-chamber). The cells were stabilized by the addition RNAprotect reagent (QIAGEN, Germantown, MD) and total RNA was purified using a RNeasy Protect Bacteria Mini Kit (QIAGEN) from three biological replicates of each strain cultured under control conditions and in exudates. The quality assessment of the extracted RNA samples was performed with a NanoDrop One^C^ Spectrophotometer (Thermo Fisher Scientific) and a 2100 Bioanalyzer (Agilent Technologies) and revealed A_260_/A_280_ and A_260_/A_230_ values of > 2.0 and a mean RNA Integrity Numbers (RIN) value of > 9.2.

Three biological replicates of RNA samples were shipped on dry ice to the DOE Joint Genome Institute (Walnut Creek, CA), where rRNA was depleted and stranded RNA-Seq libraries were prepared, quantified by qPCR and sequenced using a HiSeq 2500 instrument (Illumina). The fastq file reads were filtered and processed with BBDuk (https://sourceforge.net/projects/bbmap/) to remove reads that contained 1 or more ‘N’ bases, had an average quality score across the read less than 10 or had a minimum length < 51 bp or 33% of the full read length. Reads mapped with BBMap (https://sourceforge.net/projects/bbmap/) to masked human, cat, dog and mouse references at 93% identity were removed. Another category of removed sequences matched RNA spike-in, PhiX, common microbial contaminants and ribosomal RNAs. The processed reads from each library were aligned to the reference genome using BBMap with only unique mappings allowed (BAMs/ directory). If a read mapped to more than one location it was ignored. featureCounts (Liao et al., 2014) was used to generate raw gene counts, which were normalized to adjust for the length of each gene and total number of reads mapped for each library. The normalization formula used: n = (r/(l/1000))/(t/1000000), where n = normalized read count for gene G for library L; r = raw read count for gene G for library L; l = gene G length; and t = total reads mapped for library L. Raw gene counts were used to evaluate the level of correlation between biological samples using Pearson’s correlation.

### Bioinformatic Analysis

Count tables generated by the JGI RNA-Seq pipeline were input into DESeq2 (Love et al., 2014) to normalize and determine differential expression. Statistical significance was established through DESeq2 by using three biological replicates for control and root exudate conditions. Scatterplots were generated from the DESeq2 data table outputs using ggplot2. Genes differentially expressed between control and root exudate samples (log_2_ fold-changes −2≥ to ≤2, adjusted *p* value ≤ 0.05) were used in downstream analysis. The core genome and pangenome for the *Pseudomonas* strains used in this study were computed using the OthoMCL v.2.0, Species Tree Builder v.2.2.0, and Phylogenetic Pangenome Accumulation v1.4.0 apps implemented in the U.S. Department of Energy Systems Biology Knowledgebase (KBase) (Arkin et al., 2018). Additional comparisons were conducted with the PGAweb pan-genome analysis pipeline (Chen et al., 2018). Differentially expressed genes were assigned to core, non-core, and singleton parts of each strain’s proteome by BLASTp with an E-value cutoff of e-06, identity of 40% and coverage of 60%. Functional annotation of differentially expressed genes was carried out with the Blast2GO (Conesa and Gotz, 2008) and by manual curation using sequence comparison tools implemented in the Integrated Microbial Genomes (IMG) database (Markowitz et al., 2012), Pseudomonas Genome Database (Winsor et al., 2008), Kyoto Encyclopedia of Genes and Genomes (KEGG) (Kanehisa et al., 2008), and Geneious 10.2.3 (Biomatters, Auckland, New Zealand). Metabolic functions encoded by the differentially expressed genes were mapped using iPath 3.0 (Darzi et al., 2018). Phylogenetic analyses were carried out by building multiple sequence alignments with MAFFT v7.222 (Katoh and Standley, 2013) and inferring neighbor-joining (NJ) phylogenies with Geneious Tree Builder. The resultant phylogenetic trees were visualized with iTOL (Letunic and Bork, 2016). Reproducibility of clades within the inferred NJ trees was assessed by bootstrap re-sampling with 1,000 replicates.

### Characterization of Carbon Source Utilization with Biolog Phenotype Microarrays

The utilization of carbon sources was analyzed using Phenotype MicroArrays (Biolog, Hayward, CA, USA) as follows. The bacteria were cultured overnight on Luria-Bertani agar at 25°C, after which cells were harvested and suspended in inoculating fluid (IF-0). The transmittance of the suspension was adjusted to 42% using a Biolog turbidimeter. The cell suspension was mixed with IF-0 containing Dye Mix A (Biolog) to achieve a final transmittance of 85%. One hundred microliter aliquots of the adjusted cell suspension were inoculated into PM01 and PM02A plates, which were then incubated in an OmniLog Phenotype MicroArray System (Biolog) at 25°C for 48 h. The formation of formazan was recorded at 15 min intervals, and data were analyzed using OmniLog Parametric Analysis software v1.20.02 (Biolog). Relative growth of the studied strains was normalized to growth on D-glucose and visualized using Heatmapper (Babicki et al., 2016).

### Data Availability

Sequences generated in this project were deposited under NCBI BioProject accession numbers PRJNA439743 through PRJNA439790.

## RESULTS

### Metabolomic Profiling of Root Exudates of *B. distachyon*

Metabolomics analysis of lyophilized root exudates revealed the presence of numerous plant metabolites, 86 of which were identified by matching their spectra to the LECO/Fiehn Metabolomics library (**Table S2**). These metabolites included i) carbohydrates and their derivatives (glucose, fructose, xylose, sucrose, trehalose, maltose, galactose, and others); ii) sugar alcohols (β-mannosylglycerate, myo-inositol, galactinol, 2-deoxyerythritol, ribitol, threitol and cellobitol); iii) amino acids and derivatives (glutamine, tyrosine, glutamic acid, asparagine, aspartic acid, valine, phenylalanine, isoleucine, glycine, serine, proline, leucine, tryptophan, cysteine, methionine, citrulline, and others); iv) organic acids (aconitic, allantoic, γ-aminobutyric, azelaic, citric, fumaric, 2-furoic, D-glyceric, 3-hydroxypropionic, α-ketoadipic, malic, methylmalonic, nicotinic, quinic, succinic, threonic); and v) assorted metabolites including heterocyclic compounds, phenolics, biogenic amines, etc. (3-hydroxypyridine, maleimide, noradrenaline, 4-hydroxy-3-methoxybenzoate, 5-methoxytryptamine, uracil, aminomalonic acid, palmitic acid, urea). Results of the analysis also revealed that rhizodeposits of *B. distachyon* contain hydroxyectoine and the quaternary amine (QA) glycine betaine (**Figure S1**).

### Phylogenetic and pangenome analyses of Pseudomonas strains used in the study

We used a set of phylogenetic markers suggested by Mulet *et al*. (Mulet et al., 2010) to investigate the relatedness of the eight strains used in this study to distinct lineages recognized within the *P. fluorescens* species complex. The multilocus sequence analysis based on the concatenated sequences of the housekeeping genes *rrs* (16S rRNA), *gyrB*, *rpoB*, and *rpoD* identified R1-43-08 (along with strains 2-79 and SBW25) as a member of the *P. fluorescens* subgroup (**Figure 1**). The rest of the strains clustered closely with four additional subgroups of the *P. fluorescens* complex, namely *P. corrugata* (strains Q2-87 and Q8r1-96), *P. koreensis* (Pf0-1), *P. protegens* (Pf-5), and *P. chlororaphis* (30-84). The genomes of the eight rhizosphere *Pseudomonas* strains varied in size by 1.43 megabase (ranging from 5.65 to 7.07 Mb) and contained between 5166 and 6363 protein-coding genes (**Figure 2A**). The shared gene content was characterized with OrthoMCL, which uses all-against-all BLASTp followed by the Markov Cluster algorithm to identify protein groups shared between the compared genomes, as well as groups representing species-specific gene expansion families (Li et al., 2003). The pangenome analysis revealed a core comprised of approximately 3179 orthologs that were shared among all strains and represented 50.0% to 61.5% of each predicted proteome (**Figure 2A, 2B**). The non-core pangenome contained genes shared by two or more (but not all) strains and contained between 1482 and 2080 orthologs, which corresponded to 28.7% - 36.3% of individual proteomes. The rest of the predicted protein-coding genes were strain-specific singletons that comprised 7.5% to 15.1% of the strain’s predicted proteomes. In respect to divergence from the core genome, strain Pf-5 was found to possess the highest proportion of unique genes (*n* = 949) followed by 2-79 (*n* = 887). The entire pangenome of the *Pseudomonas* strains encompassed over 12000 homolog and singleton gene families.

**Figure 1.**
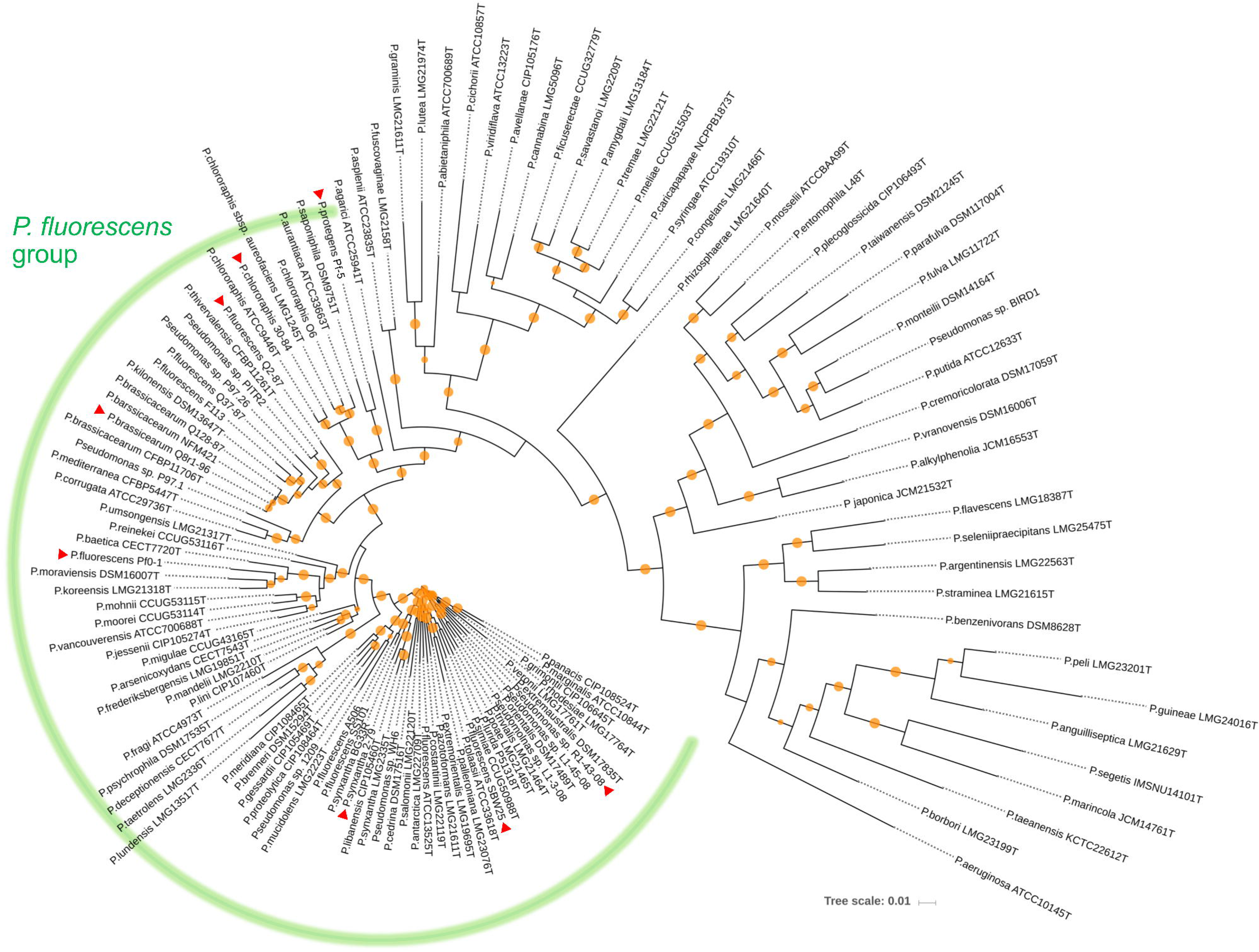
Neighbor joining phylogeny showing the relationship of the eight strains used in this study (indicated by red triangles) to different species of the *P. fluorescens* complex. The phylogeny was established based on the concatenated sequences of the housekeeping genes *rrs* (16S rRNA), *gyrB* (subunit B of DNA gyrase), *rpoB* (β subunit of RNA polymerase), and *rpoD* (sigma 70 factor subunit of RNA polymerase). Distance matrices were calculated by the Jukes-Cantor method. Colored circles on tree nodes indicate bootstrap values (1000 replicates) that vary between 60% (smallest circle) and 100% (largest circles).

**Figure 2.**
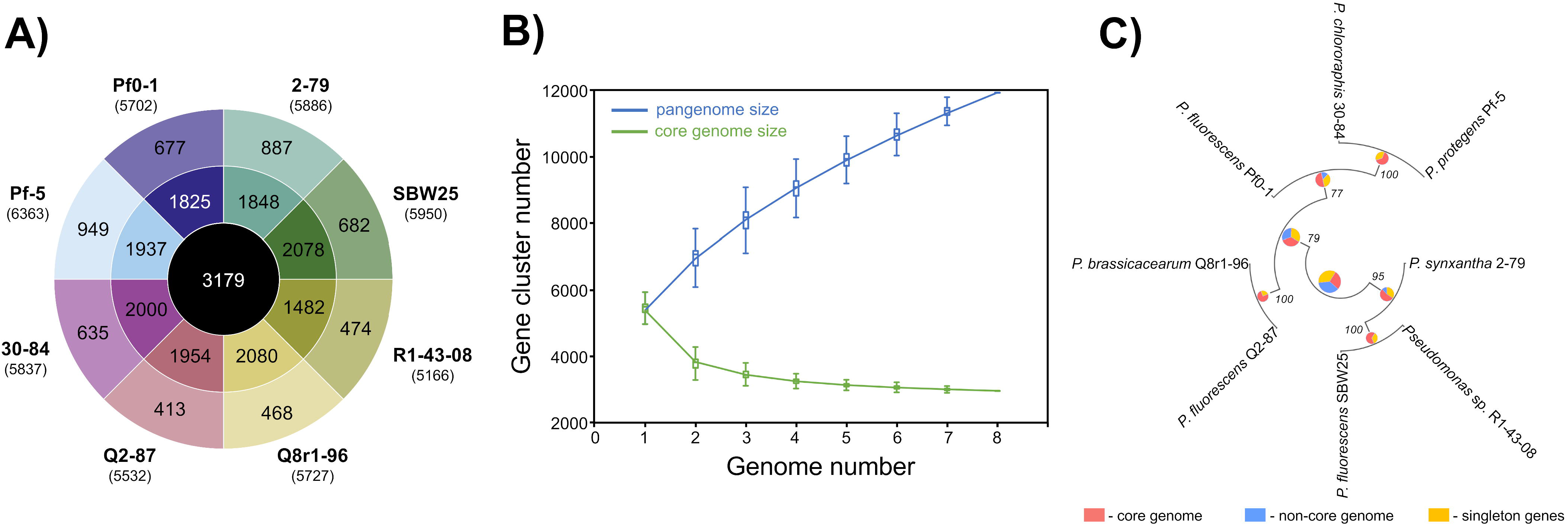
Pangenome analysis of the studied *Pseudomonas* strains. (A) The innermost circle shows the number of orthologous protein families shared among all eight strains used in this study. The second circle shows orthologs present in two or more (but not all) strains, whereas the outermost circle represents strain-specific singletons. Values in brackets under strain names correspond to the total number of protein-coding genes predicted in each genome. (B) The accumulation of gene clusters within the pan-genome and core-genome plotted against the number of genomes. (C) The partitioning of orthologs into the core, non-core, and singleton parts of the pangenome superimposed on the Maximum Likelihood genome phylogeny of the studied strains. The analysis was conducted in KBase (Arkin et al., 2018). Bootstrap support values are indicated at the nodes.

Further homolog family-based comparisons identified Q8r1-96 and R1-43-08 as the most distantly related strains, with 3349 shared homologs (**Table S3**). Q8r1-96 and Q2-87, which shared 4489 homologs, were the most closely related strains. The partitioning of homolog gene families into the core, non-core, and singleton parts of the pangenome agreed with phylogenetic relationships of the strains deduced from the analysis of a selected subset of COGs (Clusters of Orthologous Groups) (**Figure 2C**). The COG-based phylogeny supported the multilocus sequence analysis and revealed that the eight *Pseudomonas* strains form three distinct clusters, the first of which contained 2-79, R1-30-84, and SBW25. The second cluster included Q8r1-96 and Q2-87, whereas the third encompassed strains 30-84, Pf-5, and Pf0-1.

### Correlating the Composition of Root Exudates with Metabolic Profiles of *Pseudomonas* Strains

We used the Phenotype MicroArray PM1 and PM2 plates to profile the eight *Pseudomonas* strains for the utilization of 190 different carbon sources. Results of the analysis identified 90 compounds that supported growth and clustered by their intensities of utilization into three distinct groups (**Figure 3**). Group I was comprised of 30 highly metabolized carbon sources, which included several amino acids and intermediates of glycolysis, pyruvate metabolism, and citrate cycle. Approximately half of these compounds were catabolized by all eight strains, and included several organic acids (fumaric, citric, gluconic, malic, pyroglutamic), amino acids (Glu, Asn, Gln, Asp, Pro, Ala, γ-aminobutyric acid), carbohydrates (glucose, mannose, mannitol), and the purine nucleoside inosine. Group II was comprised of 44 chemically diverse carbon sources that were variably utilized by the strains. These compounds were carbohydrates, organic acids, amino acids, phenolics, and polyols, and included known compatible solutes and intermediates of metabolism of pentoses, galactose, starch and sucrose. Group III encompassed the rest of the Phenotype MicroArray test panel and contained compounds that were not catabolized by the tested strains. Among several notable exceptions were α-hydroxyglutamic acid-γ-lactone, putrescine, and itaconic, citramalic, and succinamic acids, which supported the growth of strains 2-79, 30-84, Pf-5, and SBW25. We further matched the carbon metabolic profiles of the *Pseudomonas* strains against the list of plant-derived metabolites from the rhizodeposits of *B. distachyon* Bd21. Interestingly, many carbon sources from the Phenotype MicroArray panel were also present in the root exudates of *B. distachyon* Bd21, and some of these compounds (glucose, mannose, galactose, fructose, γ-aminobutyric acid, aspartic acid, citric acid, malic acid, fumaric acid, quinic acid, alanine, glutamine, glutamic acid) were catabolized by all strains used in this study, while others (e.g., xylose, trehalose, *m*-inositol) were actively utilized only by certain organisms (**Figure 3**). The comparison of catabolic profiles across the eight studied *Pseudomonas* strains revealed the presence of three distinct clusters. The first cluster contained strains Q8r1-96 and Q2-87, which consumed very similar sets of carbon sources, as well as strain Pf0-1. The second cluster was comprised of 2-79, R1-43-08, SBW25, and 30-84, whereas the third cluster was represented by a single strain, Pf-5. The overall similarity of the catabolic profiles partially agreed with the separation of the strains into different subgroups of the *P. fluorescens* complex (see above).

**Figure 3.**
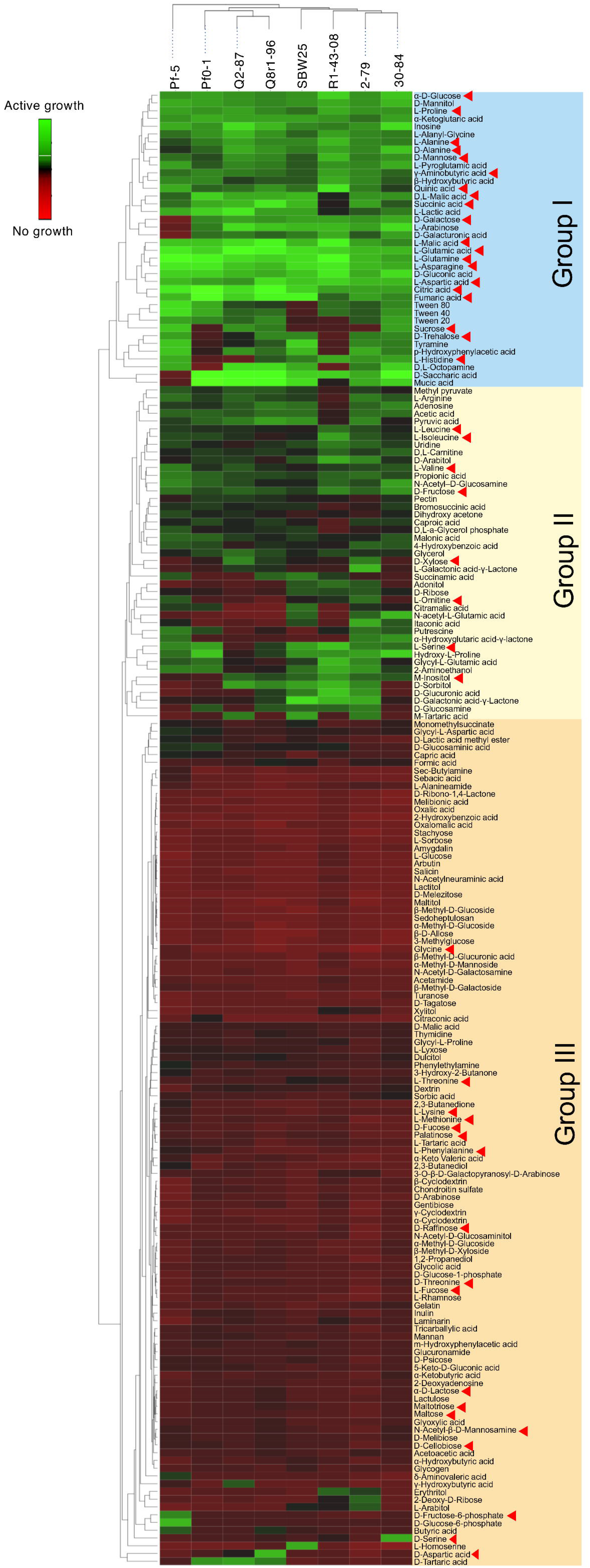
Biolog Phenotype MicroArray profiling the eight rhizosphere *Pseudomonas* strains used in the study. The hierarchical clustering analysis was carried out using the average linkage method with Euclidean distances. Carbon sources identified by red arrowheads were also detected in the sterile root exudates of *B. distachyon* Bd21.

### Analysis of the RNA-seq Results

In order to understand the cellular responses of rhizosphere *Pseudomonas* to plant exometabolites, we analyzed the transcriptome changes in cultures grown in the presence of root exudates. Under field conditions, rhizobacteria colonize plant roots in the form of surface-attached microaerobic biofilms (Hojberg et al., 1999). To mimic these conditions, the eight *Pseudomonas* strains were grown statically at 72% air saturation in 21C culture medium amended with root exudates and then processed to extract total RNA. A total of 995 million raw sequencing reads was generated from the RNA samples by using the Illumina HiSeq-2500 platform, averaging 20.7 million reads per sample. The removal of low-quality and rRNA sequences resulted in a total of 793 million filtered reads that were mapped onto the eight *Pseudomonas* genomes with a mean of 7.48 million mapped fragments per genome. The differentially abundant transcripts were identified by setting a *p* value of 0.05 (adjusted for multiple testing) and the log_2_ fold-change (FC) threshold ≥ +/−2.0 (**Figure 4; Tables S4-S11**). When compared with the control conditions, an average of 204 genes per strain were differentially expressed in the presence of root exudates, with the highest (*n* = 425) and lowest (*n* = 112) numbers observed, respectively, in SBW25 and Q2-87 (**Figure 4**). Overall, more genes were induced than repressed in response to exudates, but the actual numbers in each category varied substantially depending on the identity of the *Pseudomonas* strain. In most strains, the bulk of the differentially expressed genes was almost equally distributed between the core (mean, 48.2%) and non-core (mean, 45.8%) parts of the genome, whereas the strain-specific singleton genes constituted on average only 5.9% (**Figure 4B**). One notable exception was observed in Q8r1-96, where all differentially expressed genes belonged to the core (73.8%) and non-core (26.2%) parts of the genome. Another notable pattern was observed in R1-43-08, where the majority of genes affected by the presence of root exudate fell into the non-core category (56.3%). The highest proportion of differentially expressed singletons (11.3% and 10.4%, respectively) was identified in strains SBW25 and Pf-5.

**Figure 4.**
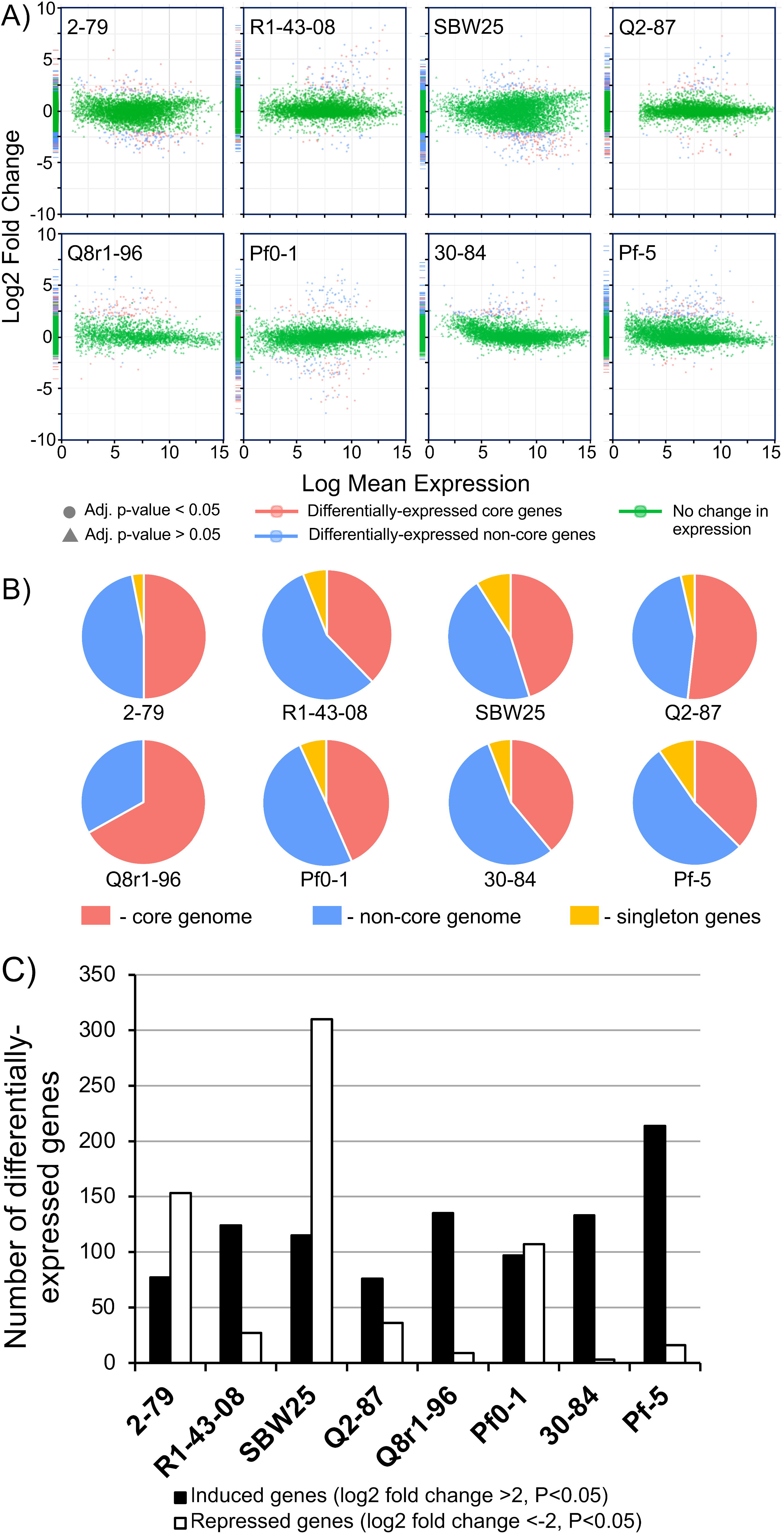
(A) Log ratio versus abundance plots (MA-plots) showing the changes in gene expression in response to root exudates. The differentially expressed core and non-core genes are shown in red and blue, respectively. Green color indicates genes with a log_2_ fold-change and/or adjusted *p* values below the established threshold. (B) Circular diagrams depicting the distribution of differentially expressed genes among the core, non-core, and singleton proteomes of individual *Pseudomonas* strains. (C) The number of genes per genome that were induced and repressed by *B. distachyon* root exudates.

We further explored how the identified differentially expressed genes were distributed across genomes of the eight studied rhizosphere strains. The pairwise BLASTp comparisons identified 2-79 and SBW25 as two strains that shared the highest number of genes (*n* = 101) induced or repressed in response to root exudates (**Table 1**). The second pair of strains with a significant number of similar differentially expressed genes (*n* = 86) was Q8r1-96 and Pf-5, which was followed by Pf0-1 and 30-84, which shared 56 differentially expressed genes. These patterns of shared genes were also observed when the results of the pairwise BLASTp comparisons were converted into a binary gene presence/absence matrix, which was then subjected to cluster analysis using a UPGMA algorithm based on Sorensen’s dissimilarity index or examined by non-metric multidimensional scaling (NMDS) (**Figure 5**).

**Table 1.**
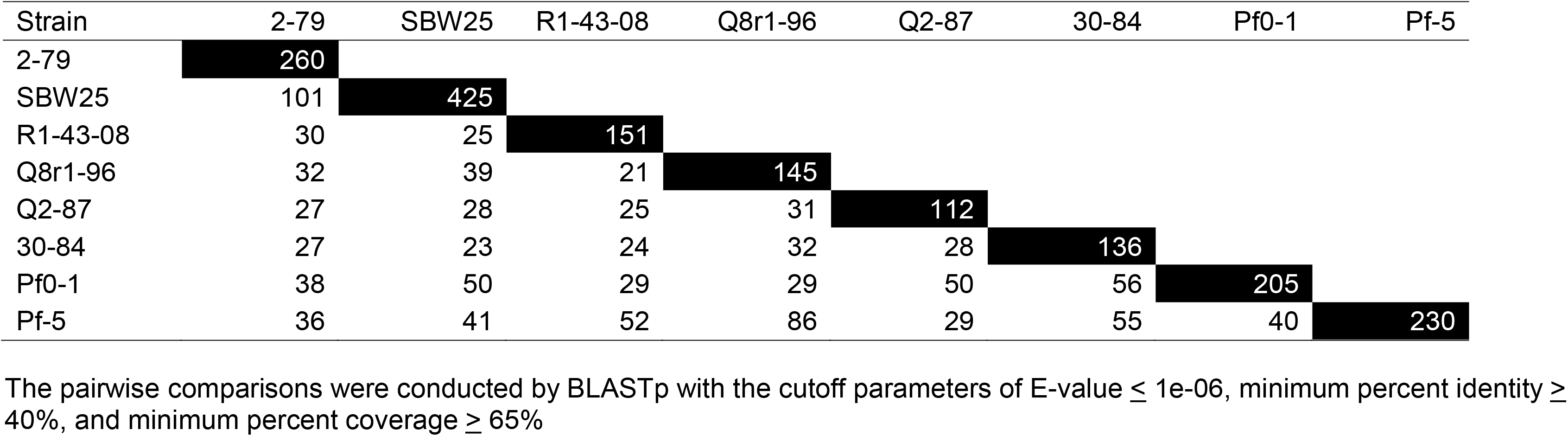
The number of differentially expressed genes shared among the eight studied strains of rhizosphere *Pseudomonas*.

**Figure 5.**
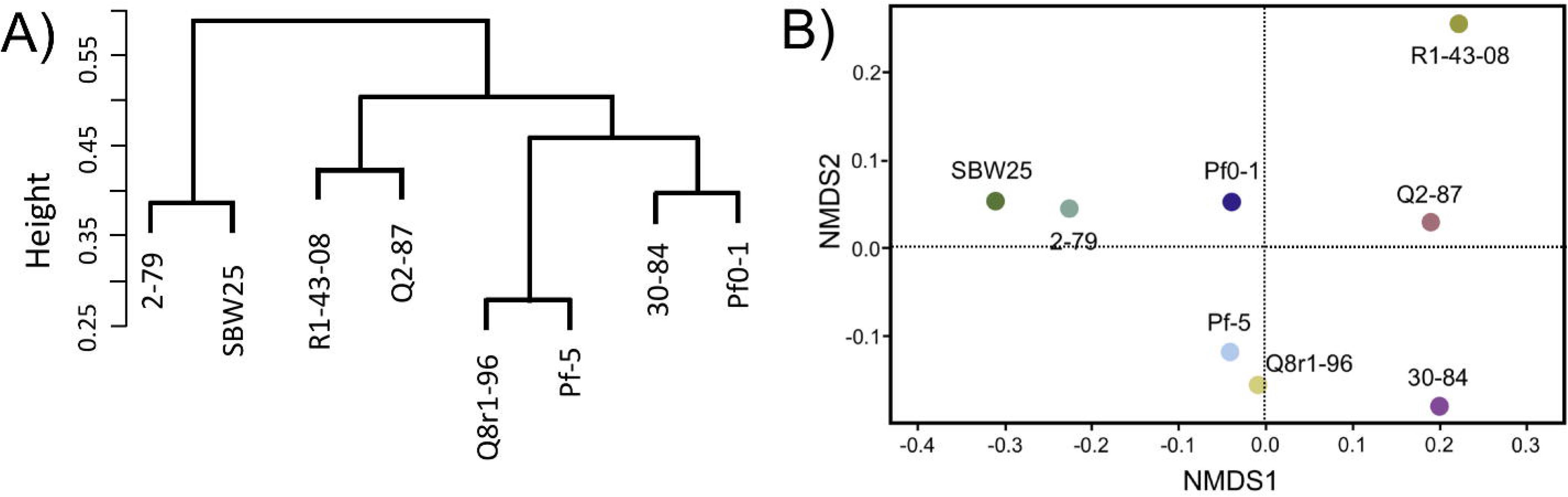
Comparison of the eight *Pseudomonas* strains based on the content (presence/absence) of genes differentially expressed in the presence of root exudates. (A) UPGMA clustering based on the Sorensen’s dissimilarity index; (B) non-metric multidimensional scaling (NMDS) analysis.

The differentially expressed *Pseudomonas* genes were subjected to Blast2Go analysis and Gene Ontology (GO) annotation (**Figure 6**). Metabolic process, catalytic activity, and membrane were the most common annotation terms across the three primary GO term categories (i.e., Biological Process, Molecular Function, and Cellular Component). A total of 1,694 GO terms was assigned to 805 upregulated genes, with the majority of the GO terms related to molecular function (682, 40.3%), followed by biological process (669, 39.5%), and cellular component (343, 20.2%). In the 539 downregulated gene category, 1,101 GO terms were assigned to biological process (420, 38.1%), molecular function (417, 37.9%), and cellular component (264, 24.0%). Within biological process, metabolic process, cellular process, localization, response to stimulus, and regulation were over-represented. Within molecular function, the largest proportion was assigned to catalytic activity, binding, and transporter activity categories. Within cellular component, the majority were assigned to membrane, membrane part, cell, and cell part categories. Across the eight strains, 37-42% of differentially expressed genes had no Gene Ontology IDs and encoded various conserved hypothetical proteins.

**Figure 6.**
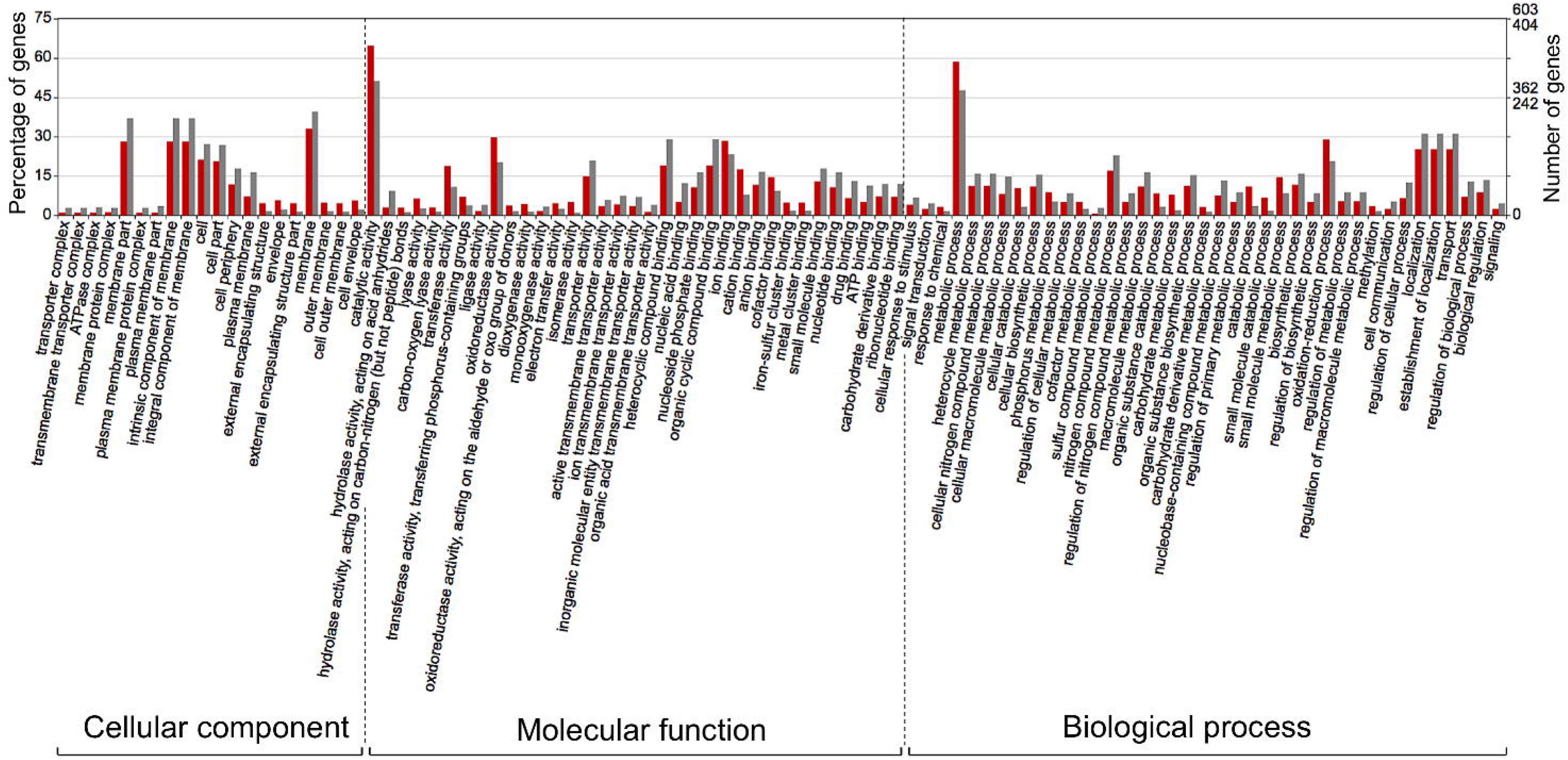
Gene Ontology (GO) classification of *Pseudomonas* genes that were induced (red bars) or repressed (grey bars) in response to root exudates of *B. distachyon* Bd21. The terms were derived from 93 different functional groups (GO sub-categories level 4).

### Functional Classification of Shared Differentially Expressed Genes

The interrogation of RNA-seq data revealed multiple cellular pathways that were differentially regulated in bacterial cultures incubated with root exudates (**Figures S2, S3**). Although none of these differentially regulated pathways were shared by all eight strains, the cross-strain comparisons revealed several types of common and specific transcriptomic responses that were elicited by the presence of plant exometabolites. The first category of shared differentially expressed pathways functioned in the uptake and catabolism of selected carbohydrates, quaternary ammonium compounds (QAs), and phenolics. All strains except for R1-43-08, responded to root exudates by inducing the fructose-specific phosphoenolpyruvate (PEP)-dependent phosphotransferase system (PTS^Fru^) (**Table 2**). The components of this system are encoded by a conserved operon and include the cytoplasmic polyprotein EI/HPr/EIIA^Fru^ (FruB), the 1-phosphofructokinase FruK, and the fructose-specific permease EIIBC (FruA) (Chavarria et al., 2016). The PTS^Fru^ system functions by acquiring high-energy phosphates from PEP and sequentially passing them, via the EI/HPr/EIIA^Fru^ domains of FruB, to the EIIB component of FruA. The phosphates are ultimately transferred by the EIIC transporter to fructose yielding fructose 1-phosphate, which is channeled into the central metabolic pathways through the action of the phosphofructokinase FruK.

**Table 2.**
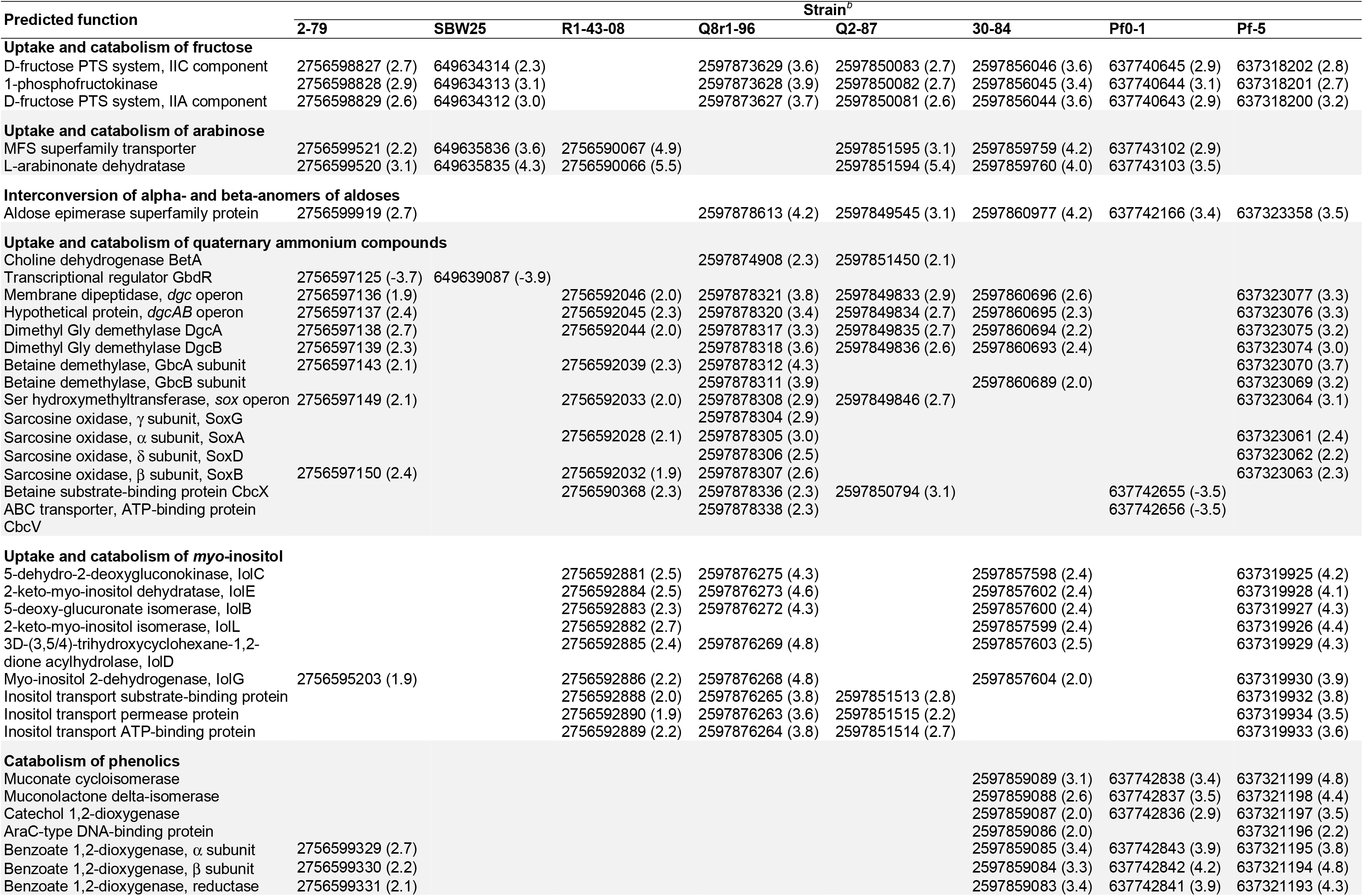
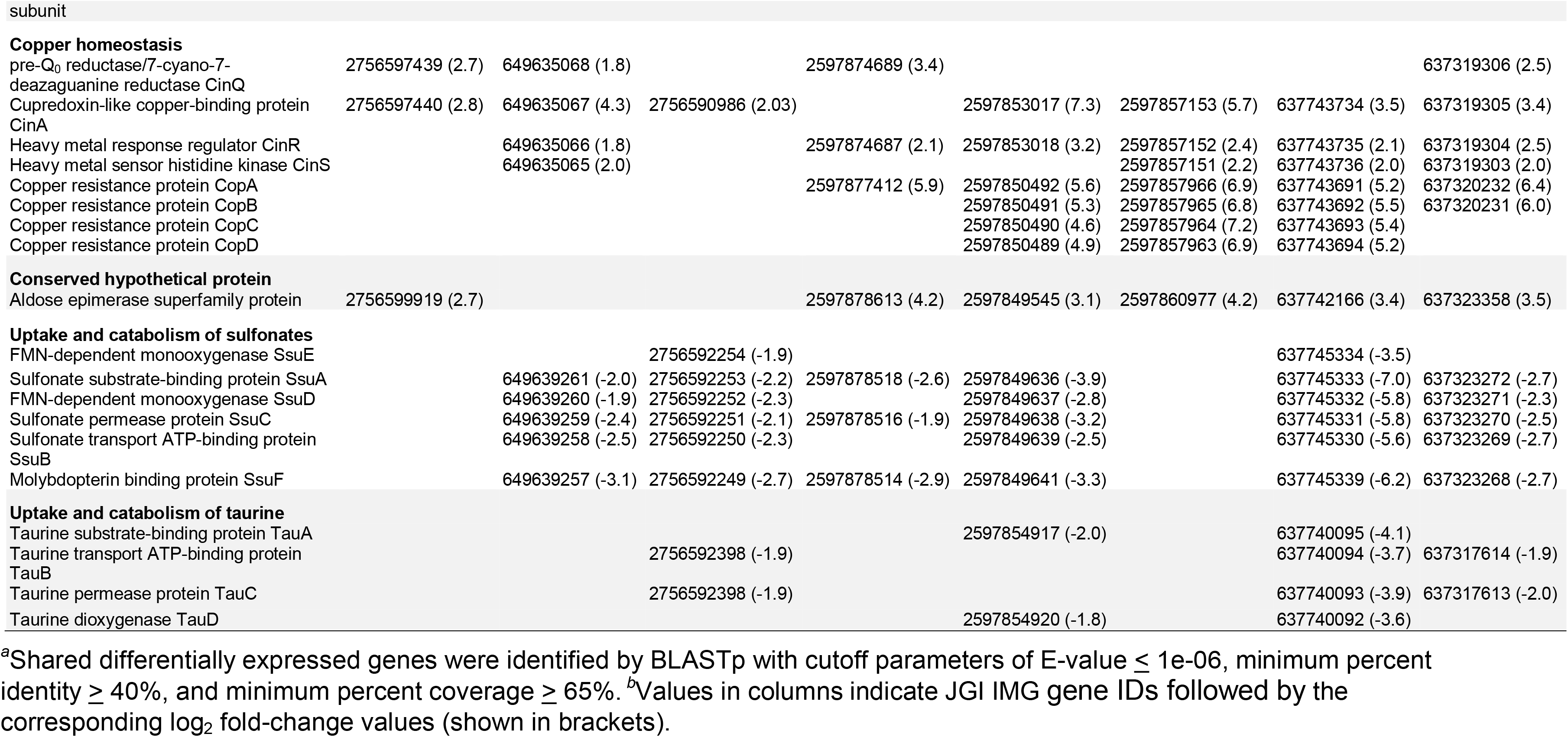
The distribution and predicted functions of selected differentially expressed genes.^*a*^

In all strains except for Q8r1-96 and Pf-5, the exposure to root exudates resulted in the induction of two genes adjacent to the *fru* cluster that encoded a Major Facilitator Superfamily (MFS) transporter and an L-arabinonate dehydratase (**Table 2**). These genes are predicted to participate in the uptake and catabolism of L-arabinose, where L-arabinonate dehydratase plays an important role by converting L-arabinonate to 2-dehydro-3-deoxy-L-arabinonate (Rahman et al., 2017). In SBW25, R1-43-08, and Q2-87 we also observed the induction of genes encoding components of the AraFGH complex, an ATP-Binding Cassette (ABC) superfamily transporter involved in the import of arabinose into the cell (**Tables S5, S6, S8**). Finally, all strains except SBW25 and R1-43-08 responded to the presence of exudates by upregulating a conserved gene encoding an aldose epimerase superfamily protein. Such enzymes equilibrate alpha- and beta-anomers of aldoses and ensure that stereospecific enzymes involved in the metabolism of free sugars do not act as metabolic bottlenecks (Abayakoon et al., 2018). Although some aldose epimerases have been linked to specific pathways, the *Pseudomonas* gene identified in this study could not be assigned to a particular metabolic process based on sequence analysis and genomic location.

Several *Pseudomonas* strains responded to the presence of root exudates by upregulating genes involved in the uptake and catabolism of *myo*-inositol and possibly other stereoisomers of inositol (**Table 2**). The upregulated catabolic genes encode the dehydrogenase IolG, which oxidizes *myo*-inositol to its corresponding ketone, as well as IolE, -D, -B, and -C that collectively convert the 2-keto-*myo*-inositol to acetyl-CoA and the glycolysis intermediate dihydroxyacetone phosphate (Yoshida et al., 2008; Kohler et al., 2011). In R1-43-08, Q8r1-96, Q2-87 and Pf-5, the upregulated functions also involved components of the putative inositol-specific ABC transporter. The cross-genome comparisons revealed that in all studied strains except for Pf0-1, components of the *myo*-inositol utilization pathway were encoded within a well-conserved gene cluster which, in addition to catabolic and transport functions, also encodes a dedicated transcriptional repressor.

All studied strains of *Pseudomonas* carry multiple genes involved in scavenging the quaternary ammonium compounds choline, glycine betaine (GB), carnitine, choline-O-sulfate, and sarcosine from the environment. Many of these genes were differentially expressed, including those encoding parts of the ABC transporter CbcXWV, which is predicted to function in the uptake choline under water-replete conditions (**Table 2**). Among enzymes induced in the presence of root exudates were the choline dehydrogenase BetA, which converts choline to glycine betaine and a network of enzymes (i.e., the Rieske family oxygenase GbcAB, the dimethyglycine demethylase DgcAB, and the sarcosine oxidase SoxBDAG) that sequentially convert GB to glycine. In 2-79 and SBW25, this group of differentially-regulated genes also included an AraC-family transcriptional activator GbdR, which perceives intercellular levels of GB and induces genes involved in the transport and catabolism of glycine betaine and detoxification of the catabolic byproducts (Hampel et al., 2014).

The last category of activated catabolic pathways included the catechol branch of the β-ketoadipate pathway for the degradation of aromatic compounds. In strains 30-84, Pf0-1, and Pf-5, growth on root exudates resulted in upregulation of catechol-1,2-dioxygenase, muconate cycloisomerase, and muconolactone isomerase, which collectively cleave the catechol ring and convert it to β-ketoadipate enol-lactone (Harwood and Parales, 1996). Finally, analysis of the *P. synxantha* 2-79 transcriptome identified an induction of *benABC* genes encoding subunits of benzoate 1,2-dioxygenase, an oxidoreductase that generates catechol from benzoate.

In addition to various catabolic pathways, the exposure to root exudates also induced several genes involved in the homeostasis of copper (**Table 2**). Four of these genes form a conserved cluster in genomes of the strains and encode the periplasmic copper-sensing two-component system CinRS, the plastocyanin/azurin-like protein CinA, and the NADPH-dependent pre-Q_0_ reductase CinQ. Also, in strains Q2-87, 30-84, Pf0-1, and Pf-5, we observed upregulation of a conserved operon encoding the multicopper oxidase CopA, the periplasmic copper-binding protein CopC, the inner membrane protein CopD, and outer membrane protein CopB. In several Gram-negative bacteria, these Cop proteins are thought to have dual functions and participate both in the uptake of essential copper as well as in the sequestration of excess copper in the periplasm and outer membrane.

The analysis of shared downregulated pathways revealed that most of the strains respond to the presence of root exudates by repressing genes involved in the uptake and catabolism of sulfur compounds (**Table 2**). In strains SBW25, R1-43-08, Q8r1-96, Q2-87, Pf0-1 and Pf-5, this response involved the *ssuEADCB* operon responsible for the utilization of alkanesulfonates as sulfur sources. The *ssu* operon is highly conserved in fluorescent pseudomonads and encodes the FMNH_2_-dependent monooxygenase SsuD and the NAD(P)H-dependent FMN reductase SsuE, which together catalyze the desulfonation of alkanesulfonates. Also, the *ssu* locus contains genes for the molybdopterin binding protein SsuF and the alkanesulfonate-specific ABC-type transporter consisting of the sulfonate substrate-binding protein SsuA, sulfonate permease protein SsuC, and sulfonate transport ATP-binding protein SsuB. Finally, in R1-43-08, Q2-87, Pf0-1, and Pf-5, growth on root exudates coincided with repression of the *tauABCD* operon, which allows these strains to utilize taurine (2-aminoethanesulfonate) as a sulfur source. The repressed *tau* genes encoded the 2-oxoglutarate-dependent taurine dioxygenase TauD and substrate-binding, ATP-binding, and permease components of the taurine-specific ABC transporter TauABC.

### Other Differentially Expressed Pathways

In addition to their effect on several shared cellular pathways, growth on root exudates resulted in the induction or repression numerous strain-specific genes. In closely related *P. synxantha* 2-79 and *P. fluorescens* SBW25, we observed differential expression of genes involved in energy metabolism, transport of amino acids, and surface attachment (**Tables S4, S5**). Other notable differentially expressed pathways included 2-79 gene clusters that encode enzymes for the catabolism of trehalose, a prophage, and toxin/antitoxin system, as well as the SBW25 operon predicted to control the synthesis of the capsular exopolysaccharide colonic acid. The response of *Pseudomonas* sp. R1-43-08 to root exudates also involved differential expression of different energy metabolism pathways. In addition, we observed the upregulation of genes involved in the uptake and catabolism of xylose (also upregulated in 2-79) and repression of enzymes for the biosynthesis of phenazine-1-carboxylic acid and assimilation of inorganic sulfur and L-cysteine biosynthesis (**Table S6**).

The analysis of the Q8r1-96 transcriptome revealed perturbation of different metabolic pathways including genes encoding components of cytochrome C oxidase, transport and catabolism of sorbitol/mannitol, metabolism of butanoic acid, and biosynthesis of exopolysaccharides alginate and poly-β-1-6-*N*-acetylglucosamine (**Table S7**). In *P. fluorescens* Q2-87, we identified differential expression of genes involved in metabolism of galactose, tryptophan, tyrosine, glycine, serine, and threonine (**Table S8**), while in *P. chlororaphis* 30-84, growth on exudates activated the biosynthesis of molybdopterin cofactor, catabolism of - galactonate and acetoin, and uptake and catabolism of putrescine (**Table S9**). The response of *P. protegens* Pf-5 to root exudates involved upregulation of acetoin dehydrogenase, which converts acetoin to acetaldehyde and acetyl-CoA, as well pathways for the utilization of glycolate and putrescine (**Table S11**). Also induced were genes for the production of pyrrolnitrin and PhlG hydrolase, which modulates the metabolic loads attributed to the synthesis of 2,4-diacetylphloroglucinol. The differentially expressed genes of *P. fluorescens* Pf0-1 included, among others, operons encoding cytochrome C oxidase and enzymes for catabolism of malonic acid (**Table S10**). Yet another interesting finding involved the induction of assorted genes acting in the homeostasis of iron and defense against reactive oxygen species (ROS). We observed activation of iron dicitrate transporters (SBW25 and 30-84), genes for the biosynthesis of siderophores ornicorrugatin (SBW25) and pyochelin (Pf-5), heme-degrading enzymes (2-79, 30-84), TonB siderophore receptors, and components of the energy-transducing inner membrane complex TonB-ExbB-ExbD (2-79 and Pf-5). The differentially expressed ROS defense pathways were represented by different catalases in strains 2-79, R1-43-08, Q8r1-96, Q2-87, Pf0-1, and Pf-5, and organic hydroperoxide resistance proteins in strains SBW25 and R1-43-08. Finally, in SBW25, Q2-87, 30-84, and Pf0-1, the addition of exudates resulted in the upregulation of peroxiredoxins that detoxify H_2_O_2_, peroxynitrite, and aliphatic and aromatic hydroperoxides.

## DISCUSSION

The ability of the plant microbiome to positively influence plant development, vigor, health, and fitness in response to biotic and abiotic stressors is documented by numerous studies, but molecular details of this process are largely unresolved. Complete and draft genome sequences are now available for multiple strains and species within the *P. fluorescens* group, while pathways involved in the rhizosphere lifestyle of these important microorganisms are only beginning to be systematically defined. This study contributes to such efforts by comparing transcriptomic responses to root exudates in eight well-characterized *Pseudomonas* strains that have been selected from the soil or plant rhizosphere based upon their biological control properties (**Table S1**).

Our analysis of *B. distachyon* rhizodeposits revealed a complex mix of primary and secondary metabolites, thus supporting the view of the plant rhizosphere as a carbon-rich niche for soil microorganisms. Our results were in agreement with a recent report of 27 different sugars, amino acids, and organic acids in *Brachypodium* exudates (Kawasaki et al., 2016). We confirmed the presence of exometabolites identified in that study, along with dozens of additional analytes that were identified by matching their mass-spectra and retention indices to the LECO/Fiehn Metabolomics library (**Table S2**). The complementation of the metabolomic analysis with profiling of the bacteria by Biolog Phenotype MicroArrays revealed that a substantial proportion of the characterized exudate constituents were catabolized by a collection of eight *Pseudomonas* strains from across the *P. fluorescens* group that are known to form associations with plant roots. The amendment of *Pseudomonas* cultures with root exudates caused changes in the expression of multiple genes encoding catabolic and anabolic enzymes, predicted transporters, transcriptional regulators, stress response, and conserved hypothetical proteins. In most strains, these differentially expressed genes were almost equally split between the core and variable genome regions, mirroring the substantial strain-to-strain variation in the genome size and gene content within the *P. fluorescens* species complex (Loper et al., 2012).

The analysis of transcriptome responses to root exudates revealed several types of cellular pathways present in the strains used in this study. The first category of such pathways was involved in the catabolism of carbohydrates such as fructose, arabinose, *myo*-inositol, xylose, trehalose, and galactose. Among these catabolic traits, the ability to utilize fructose as a carbon source is highly conserved among fluorescent pseudomonads. In contrast, growth on arabinose, *myo*-inositol, xylose, and trehalose is variably present and was traditionally used to differentiate species and biovars within the *P. fluorescens* group (Barrett et al., 1986). We speculate that such variably distributed pathways contribute to the differential affinity of pseudomonads towards host plants and/or to determine which strains flourish in response to growing roots and changing environments. Several independent studies have confirmed the importance of carbohydrate catabolism pathways for the biology of rhizosphere pseudomonads. For example, *in vivo* expression technology (IVET) profiling of *P. fluorescens* SBW25 identified xylose isomerase among genome regions essential for the colonization of sugar beet seedlings (Liu et al., 2015), whereas a genome-wide Tn-Seq screen of *Pseudomonas simiae* identified genes for the catabolism of *myo*-inositol among traits essential for the colonization of *Arabidopsis thaliana* roots (Cole et al., 2017).

The response of rhizosphere *Pseudomonas* to *Brachypodium* root exudates also involved pathways for the uptake and metabolism of amino acids. We observed differential expression of genes encoding the hydrophobic (HAAT) and polar (PAAT) amino acid uptake transporters in strains 2-79, SBW25, Q2-87, Pf0-1, and Pf-5. Other related genes encoded enzymes for the catabolism of valine and glutamic acid (2-79); metabolism of tryptophan, glycine, serine, and threonine (Q2-87); and biosynthesis of methionine (Q8r1-96). It is plausible that the abundance of amino acids in root exudates is also linked to the repression of pathways involved in the catabolism of sulfonates and taurine that was observed in several strains (**Table 2**). Although the preferred source of sulfur for *P. fluorescens* is unknown, in the closely related *P. aeruginosa*, the sulfur starvation response is triggered by the growth on any sulfur compound other than sulfate, thiocyanate, and cysteine (Hummerjohann et al., 1998). This fact, together with the presence of cysteine and cystine in the root exudates, suggest that rhizodeposits of *Brachypodium* may serve as an important source of sulfur for rhizosphere *Pseudomonas*. These findings also agree well with the reported scarcity of inorganic sulfate in the soil, and the presence of sulfur mostly in the form of organic compounds, including amino acids, proteins, sulfate esters, and sulfonates (Autry and Fitzgerald, 1990).

In addition to primary metabolites, plant root exudates contain simple phenolic compounds, some of which have been implicated in allelopathy and allelobiosis in wheat, barley, and rice, which are closely related to *Brachypodium* (Bertin et al., 2003). Such metabolites exhibit concentration-dependent stimulatory or inhibitory effects on plants, pathogens, and soil microorganisms (Zhou and Wu, 2012; Zwetsloot et al., 2018; Bouhaouel et al., 2019). Phenolic allelochemicals are often synthesized in response to abiotic and biotic stress, and their accumulation results in phytotoxicity and increased susceptibility to diseases (Weir et al., 2004; Tian et al., 2019). Treatment of plants with microorganisms capable of degrading such allelochemicals has been associated with phytostimulatory effects and suppression of soilborne pathogens (Chen et al., 2011). Our results revealed that several *Pseudomonas* strains responded to root exudates by inducing pathways for the catabolism of aromatic compounds, suggesting their ability to modulate the levels of phenolic allelochemicals in the rhizosphere. We speculate that the ability to degrade plant-derived phenolics may contribute to plant growth promotion and biocontrol properties of these beneficial rhizobacteria. Plant roots are not the only source of phenolic compounds, as these metabolites also enter the soil as intermediates of the breakdown of lignin (Brink et al., 2019) and polyaromatic hydrocarbons (Harwood and Parales, 1996). Therefore, the induction of aromatic catabolism in *Pseudomonas* by root exudates may have additional beneficial effects linked to stimulating the turnover of soil organic matter and mineralization of man-made soil pollutants.

Another interesting result of this study was the concerted activation of copper and iron homeostasis pathways observed in all of the *Pseudomonas* strains used in this work. In bacteria, an excess of copper is toxic and triggers oxidative stress due to the formation of free radicals, as well as disruption of protein metalation and stability of iron-sulfur clusters (Bondarczuk and Piotrowska-Seget, 2013). On the other hand, copper is an essential trace element used as a cofactor in different enzymes. Similarly, although elevated levels of iron cause redox stress, this element is also found in active energy metabolism enzymes and is crucial for bacterial growth (Andrews et al., 2003). The analysis of metal homeostasis genes identified in this study suggests that their induction was likely triggered by the deficiency of copper and iron in bacterial cultures grown in the presence of root exudates. We attribute this effect to the ability of some components of root exudates to chelate soil metals.

Despite the abundance of iron in the soil, its bioavailability is limited due to the low solubility of Fe(III) oxyhydrates at neutral pH. The non-graminaceous plants circumvent this problem by acidifying the rhizosphere and secreting flavins, phenolics, and organic acids that chelate iron. The reduction of these ferric chelates releases soluble ferrous iron taken up by root cells (Kobayashi and Nishizawa, 2012). Graminaceous plants, like *Brachypodium*, acquire iron by secreting into the soil non-protein amino acids of the mugineic acid (MA) group, which act as Fe(III)-chelating phytosiderophores. In addition to iron, low-molecular-weight organic acids and phytosiderophores bind other divalent and trivalent metals (including copper) and contribute to heavy-metal tolerance in plants (Chen et al., 2017). It is plausible that the presence of these plant exometabolites is responsible for the deficit of iron and copper observed in *Pseudomonas* cultures grown in the presence of root exudates. These results further underscore the importance of diverse and redundant metal homeostasis pathways found in genomes of the *P. fluorescens* group for the ability of these organisms to colonize and persist in the plant rhizosphere.

Recently, Klonowska and colleagues (Klonowska et al., 2018) examined transcriptomic responses of symbiotic nitrogen-fixing bacteria to root exudates of the legume plant *Mimosa pundica*, which has an unusual ability to support both alpha- (*Rhizobium*) and beta-rhizobia (*Cupriavidus* and *Burkholderia*). Using RNA-seq, the authors characterized genes involved in the perception of root exudates in the nodulating bacteria *Burkholderia phymatum* STM815, *Cupriavidus taiwanensis* LMG19424, and *Rhizobium mesoamericanum* STM3625. Interestingly, the analysis of differentially expressed genes revealed induction of pathways involved in the catabolism of fructose, xylose, *myo*-inositol, and protocatechuate/catechol. Also upregulated were some copper homeostasis, siderophore biosynthesis, and oxidative stress genes. Finally, the analytical profiling of *M. pundica* exudates revealed an overlap with rhizodeposits of *Brachypodium* in the types of carbohydrates, amino acids and organic acids present. These findings suggest that differentially expressed genes shared by multiple strains of the group *P. fluorescens* are not unique to the *Brachypodium-Pseudomonas* system but represent a set of conserved cellular pathways involved in the perception of plant exometabolites by different clades of rhizosphere-dwelling *Proteobacteria*.

Most strains included in this study were originally selected based on the ability to colonize the rhizosphere and produce secondary metabolites that alleviate the plant stress response and/or inhibit soilborne pathogens. It has been suggested that plant metabolites released into the rhizosphere affect the biocontrol activity of plant-beneficial pseudomonads (de Werra et al., 2011). We provide further support to this hypothesis by demonstrating that in some strains, root exudates modulate the expression of genes for the catabolism of the plant growth-promoting metabolites acetoin and 2,3-butanediol. The exposure to exudates also affected the expression of genes for the synthesis of well-characterized antifungal compounds pyrrolnitrin, phenazine-1-carboxylic acid, and 2,4-diacetylphloroglucinol. The modulatory effects were strain-specific, suggesting significant differences in the regulatory networks involved in the perception of plant signals and regulation of the production of antibiotics and growth-promoting metabolites.

The final significant finding of this study was the induction of catabolism of quaternary amines (QAs) observed in multiple strains of the *P. fluorescens* group during growth on root exudates. This observation was supported by the detection of glycine betaine in the root secretions of *B. distachyon*. The presence of QAs in plant tissues and the capacity of these metabolites to provide stress protection and nutrients to plant pathogens and symbionts were reported before (Boncompagni et al., 1999; Chen et al., 2013; Kabbadj et al., 2017), but our study is among the first to highlight the potential importance of these metabolites for rhizosphere interactions. Pseudomonads do not synthesize QAs *de novo* but have evolved many pathways to scavenge them from eukaryotic hosts, where these metabolites are abundant due to the prominence of phosphatidylcholine in cellular membranes. Strains of *P. fluorescens* carry genes for the conversion of choline, carnitine, and glycine betaine to glycine, as well as quaternary amine transporters of the BCCT and ABC families that are also conserved in the opportunistic human pathogen *P. aeruginosa* and the plant pathogen *P. syringae* (Galvao et al., 2006; Chen et al., 2013; Wargo, 2013b). In *P. aeruginosa*, choline catabolism genes are essential for the ability of this pathogen to persist during lung infection (Wargo, 2013a). Similarly, a *P. syringae* mutant deficient in BetT, OpuC, and CbcXWV quaternary amine transporters had reduced fitness during colonization of bean and soybean leaves under greenhouse and field conditions (Chen et al., 2013). Depending on water availability, *P. aeruginosa* and *P. syringae* catabolize exogenously supplied QAs as carbon and nitrogen sources or accumulate them as osmoprotectants (Chen et al., 2013; Wargo, 2013b). Our ongoing work in *P. synxantha* 2-79 unraveled similar physiological responses and demonstrated that QA transporters function differentially and redundantly in the uptake of quaternary amines as nutrients (Pablo, C. and Mavrodi, D., unpublished). In contrast, under water stress, the QAs choline, betaine, and carnitine are accumulated preferentially for osmoprotection. Under drought stress, a 2-79 mutant devoid of all known QA transporters was less competitive in the colonization of the *Brachypodium* rhizosphere than its wild-type parental strain. Interestingly, our metabolomic profiling of root exudates also revealed proline, glutamine, and hydroxyectoine. These metabolites act as compatible solutes in different groups of microorganisms (Yancey et al., 1982; Empadinhas and da Costa, 2008), suggesting an important role of rhizodeposits in the ability of *Pseudomonas* to persist in the rhizosphere of drought-stressed plants.

## Author contributions

DM, OM, and LT: conceived the research project; OM and JM: collected root exudates; OM and DM: cultured strains and extracted total RNA; AB and DG: performed metabolomic analysis of root exudates; DM, JP, and AF: analyzed RNA-seq data; LE, KH, and IP: conducted Biolog analyses; DM, AF, OM, DW, and LT: wrote the manuscript and all authors contributed to the manuscript revision.

