## Supplementary material for "The effect of root exudates on the transcriptome of rhizosphere *Pseudomonas* spp": Figure S1

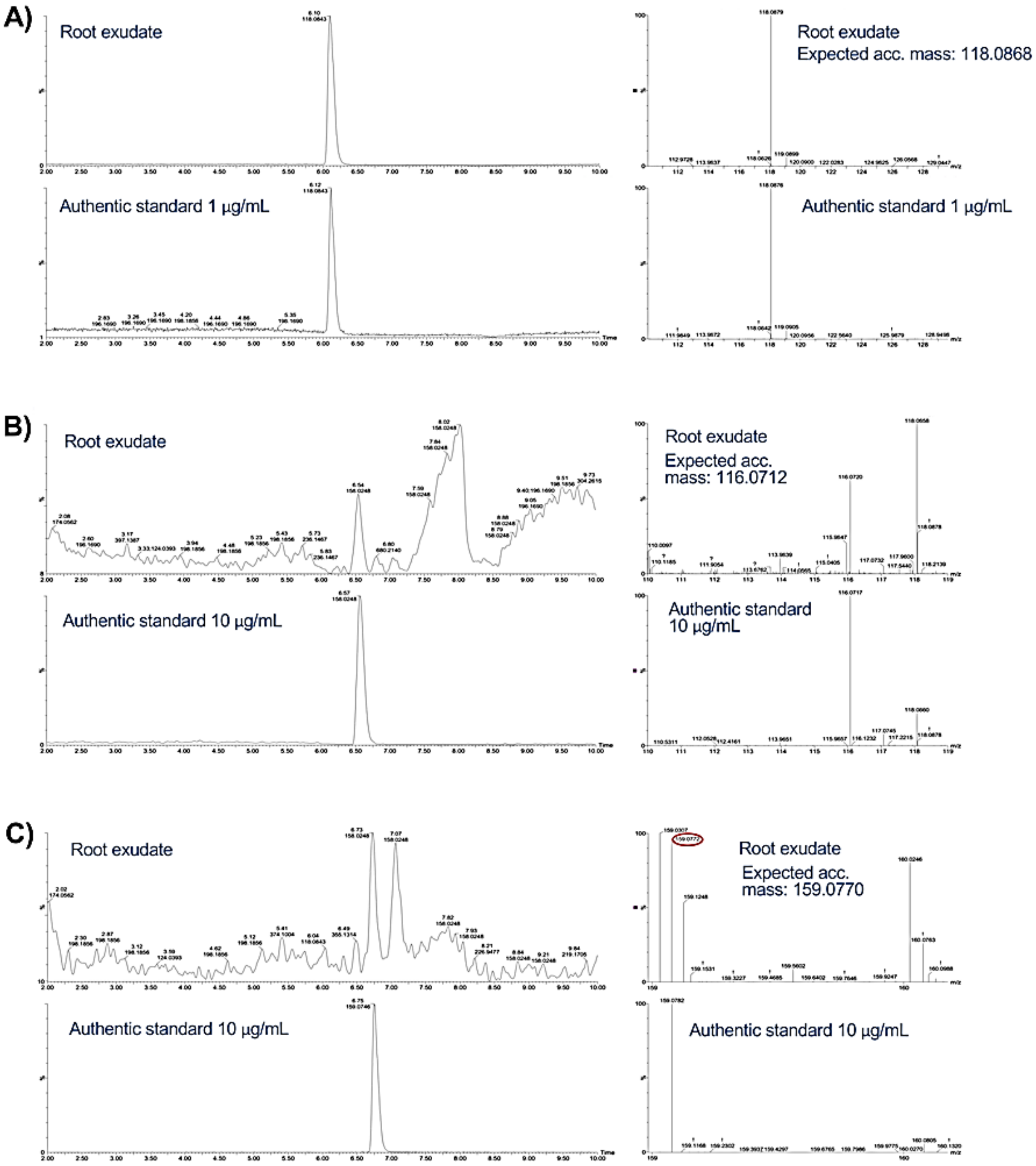


**Figure S1.** Mass chromatograms of root secretions and authentic standards showing the presence of glycine betaine (A), proline (B), and hydroxyectoine (C) in exudates of hydroponically grown *Brachypodium* plants. For each compound, panels on the left show selected ion chromatograms, while panels on the right depict accurate mass comparisons.
