## Supplementary figures and images for "The effect of root exudates on the transcriptome of rhizosphere *Pseudomonas* spp"

### Figure S2

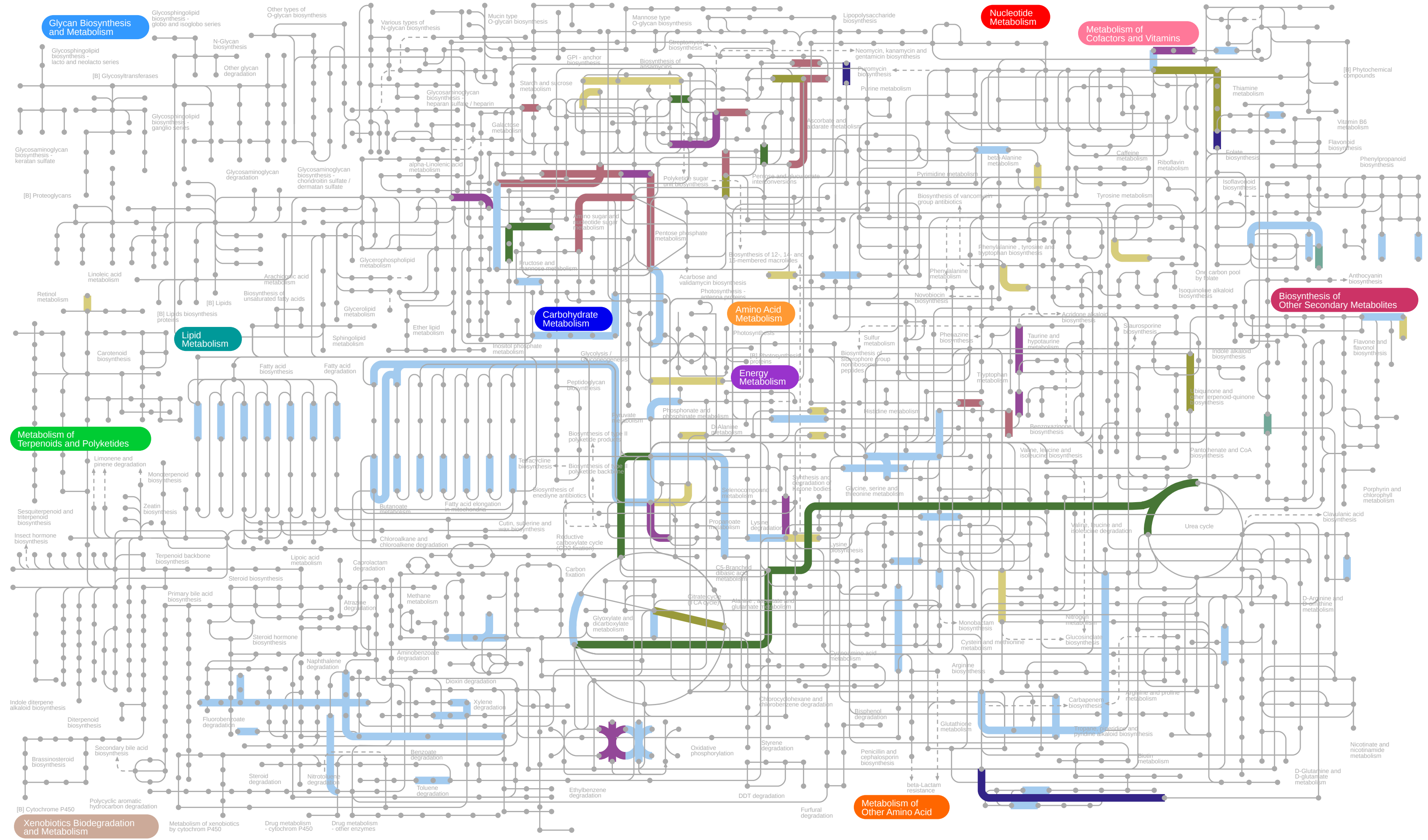


01100 5/11/17
(c) Kanehisa Laboratories

### Figure S3

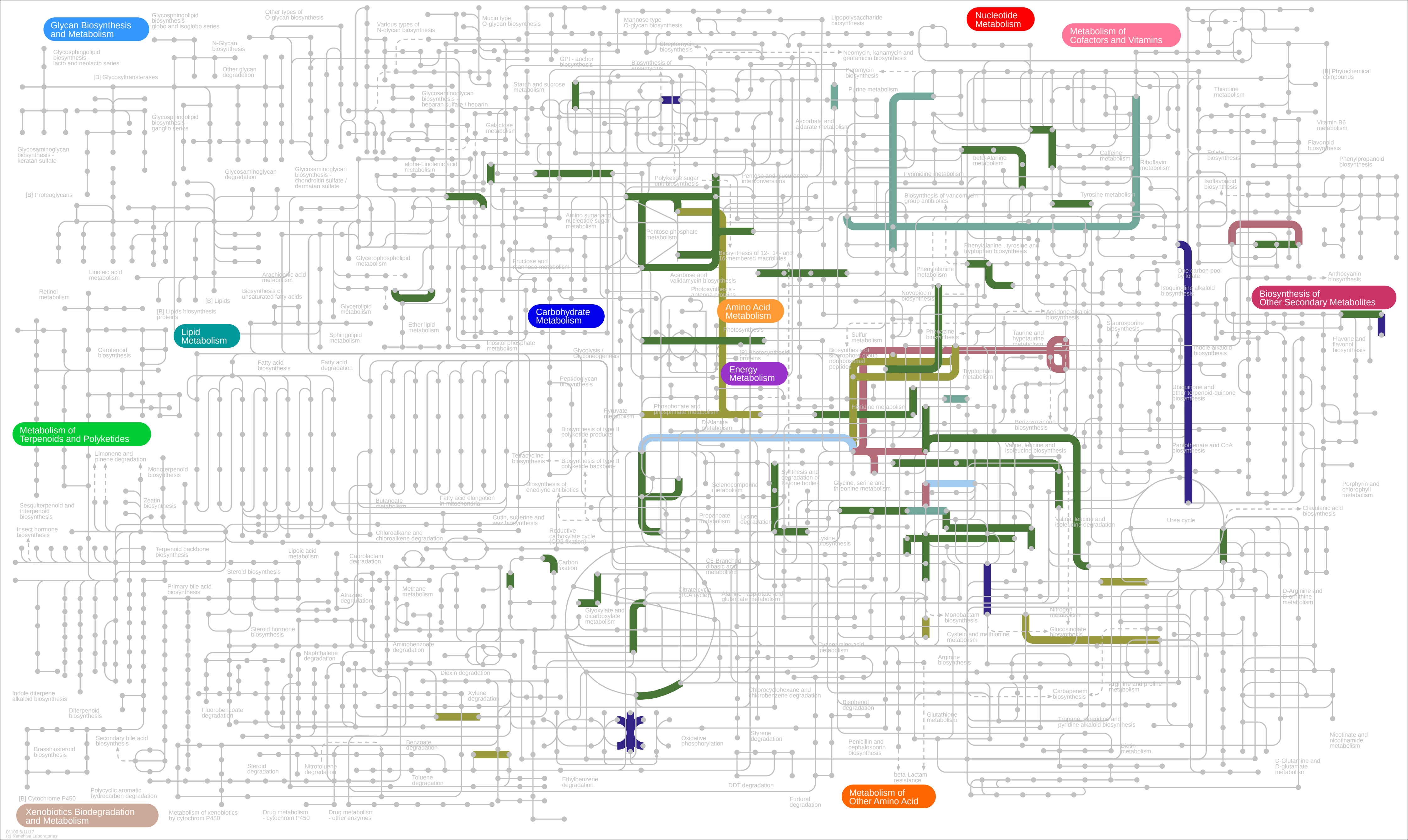
