## Supplementary material for "The effect of root exudates on the transcriptome of rhizosphere *Pseudomonas* spp": Table S1

**Table S1.** Origin and relevant phenotypes of bacterial strains used in the study.

| **Strain*^a^*** | **Origin** | **Relevant properties** | **Reference** | **Genome acc #** |
| --- | --- | --- | --- | --- |
| *Pseudomonas brassicacearum* | |  |  |  |
| Q8r1-96 | Wheat, Washington State, USA | Biocontrol of take-all of wheat caused by *Gaeumannomyces graminis* var. *tritici* | (1) | NZ_CM001512 |
| *Pseudomonas chlororaphis* sbsp. *aureofaciens* | |  |  |  |
| 30-84 | Wheat, Kansas, USA | Biocontrol of take-all of wheat caused by *G. graminis* var. *tritici* | (2) | NZ_CM001559 |
| *Pseudomonas fluorescens* | |  |  |  |
| Pf0-1 | Soil, Massachusetts, USA |  | (3) | NC_007492 |
| Q2-87 | Wheat, Washington State, USA | Biocontrol of take-all of wheat caused by *G. graminis* var. *tritici* | (4) | NZ_CM001558 |
| SBW25 | Sugar beet, Oxfordshire, UK | Biocontrol of seedling emergence disease caused by *Pythium ultimum* | (3) | NC_012660 |
| *Pseudomonas synxantha* | |  |  |  |
| 2-79 | Wheat, Washington State, USA | Biocontrol of take-all of wheat caused by *G. graminis* var. *tritici*; biocontrol of root rot of wheat caused by *Rhizoctonia solani* | (5) | CP027755 |
| *Pseudomonas protegens* | |  |  |  |
| Pf-5 | Soil, Texas, USA | Biocontrol of seedling emergence disease caused by *P. ultimum* | (6) | CP000076 |
| *Pseudomonas* sp. | |  |  |  |
| R1-43-08 | Wheat, Washington State, USA | Biocontrol of root rot of wheat caused by *R. solani* | (7) | CP027734 |

*^a^* In previous publications, strains Q8r1-96, 2-79, and Pf-5 have been designated as *P. fluorescens* Q8r1-96, *P. fluorescens* 2-79, and *P. fluorescens* Pf-5.
